# PyPlaque: an Open-source Python Package for Phenotypic Analysis of Virus Plaque Assays

**DOI:** 10.1101/2024.08.07.603274

**Authors:** Trina De, Vardan Andriasyan, Artur Yakimovich

## Abstract

Virological plaque assays are the primary method for quantifying infectious particles in a suspension, achieved by incubating a serial dilution of the virus with a monolayer of indicator cells. Existing software tools for quantification of plaque assay images lack modularity, show measurements disagreements or are closed-source - a common hurdle in BioImage analysis. We introduce PyPlaque, an open-source Python package focusing on flexibility and modularity rather than a bulky graphic user interface. Unlike previous methods, an abstracted architecture using object-oriented programming allows accommodation of various experimental containers and specimen carriers as data structures while focusing on phenotype-specific information. Aligned with the logical flow of experimental design and desired quantifications, it delivers insights at multiple granularity levels, facilitating detailed analysis. We demonstrate how this approach allows to focus on alleviating the disagreement in measurements. Furthermore, similar design is generalisable to diverse datasets in various biological contexts that fit our structural paradigm.

## Introduction

The ongoing SARS-CoV-2 pandemic proved that our ability to detect and quantify viruses is vital for stepping up our efforts to both diagnose and research emerging pathogens in a timely fashion. Yet, while a number of novel assays have been developed or improved (1–3), the virological plaque assay (4, 5) remains the only existing way to measure infectious virus isolates and quantify viral spread in tissue culture (6–8). Unlike other assays aiming to quantify a proxy of infection like viral proteins (antigen), gene products (PCR or rtPCR) or disease manifestations (chest X-Ray), the virological plaque assay measures the ability of the isolated or cultured infectious virus to infect cells and produce progeny infectious virions infecting the cells thereafter - a property remaining at the very core of virus virulence.

While slight variations of the plaque assay exist depending on the pathogen, tissue culture, format, or recording method, at its core, every plaque assay requires two main components, a highly diluted (low multiplicity of infection) inoculum and cultured indicator cells (4, 5). An infection assay set up in such a way will result in only a few infected cells, typically, within a confluent monolayer of indicator cells (4). Upon incubation for the duration of at least one full replication cycle of the pathogen, the initially infected cells will release their infectious progeny. These will, in turn, infect the neighbouring indicator cells leading to the emergence of the clonal patterns of spread. At the right dilution of the inoculum, these patterns will not overlap, allowing the observer to simply count them, thus quantifying the number of infectious particles in the inoculum measured as the number of plaque-forming units (PFU) per unit of volume. As the pathogen spreading patterns may often be intricate, measurements are often performed manually by a trained specialist.

Beyond the quantification of infectious particles, however, virological plaque assay represents a model system allowing to study virus spread phenotypes (9–11). Specifically, the size, shape, and consistency of plaques, as well as the patterns and motion trajectories of the infected cells constituting the virological plaque may reveal a great deal of information about the underlying biological mechanisms leading to the formation of such phenotypes (10–12). Therefore, harnessing this information through computational analysis of biomedical images is an invaluable tool for analysing known and emerging viruses.

To address this, apart from custom project-related or closed-source code, several approaches have been proposed (9, 11, 13–18). However, while useful for a broad range of applications the vast majority of these tools remain highly specialised. For example, Plaque 2.0 (9, 11), Infection Counter (14), and SCFQ (16) are aimed exclusively at fluorescence microscopy. At the same time, ViralPlaque (13) plugin for ImageJ/Fiji (19) software is highly tuned for well-separated crystal-violet-stained plaques. Additionally, Plaque 2.0 (9) (11), and SCFQ (16) specialise in high-content images. Plaque Size Tool (17), in turn, while more versatile from the image source perspective requires only well-separated plaques. Other tools, like Viridot (15) are dedicated to one specific virus only. Finally, all tools existing thus far, are highly reliant on user input via the graphic user interface and don’t support modern open-source data analysis languages like Python.

Such a highly scattered landscape of tools indicates that despite the seeming similarity, all these tools were developed for their own purpose. Being a fundamental assay in Infection Biology, plaque assay is performed slightly differently depending on the pathogen, imaging techniques, indicator cell types, throughput requirements, and reagent availability. Furthermore, should the purpose of the assay lay beyond the quantification of PFU, more often than not the plaque assay would be specialised to the final requirements of the study. Evidently, should a unified solution exist in such a case, it would require a great degree of flexibility while taking care of the complexity of plaque assay.

To address this, here we propose PyPlaque (20) - an open-source Python package allowing to simplify plaque assay analysis. Unlike the graphic-user-interface-based tools mentioned above, PyPlaque takes a radically different approach. Instead of a no-code solution, we provide a convenient library that lowers the amount of code required to analyse plaque assay data while maintaining ultimate analysis flexibility. We achieve this by providing Python data structures (akin to Pandas (21) for tabular data). Using these data structures, PyPlaque allows users to account for experiments, dishes or plates and plaque phenotypes. Utilising Python features like IPython/Jupyter (22, 23) interactive notebooks, such scripts or notebooks could be rapidly created and shared. PyPlaque is aimed at programming-savvy biologists and significantly simplifies the development of highly-optimised plaque assay quantification scripts. Leveraging the vast data analysis package ecosystem of Python, notebooks can be adapted by fellow researchers, who may need a set of specialised modifications. We demonstrate the versatility of our approach on several examples, including different variations of plaque staining, and imaging. Furthermore, our examples employ diverse viruses. Lastly, PyPlaque is open-source and requires only open-source software to run.

## Results

### PyPlaque Design Principles

To develop a highly customisable computational tool for virological plaque analysis we followed several design choices. Firstly, we aimed for the tool to be based upon modern and widespread open-source programming language in order to fit seamlessly into the tool-set of Data Science. This led to Python being the language of choice. Secondly, we aimed to use object-oriented programming (OOP) to create logic and structures allowing us to seamlessly capture the experiment and specimen details. We set out to create structures as seamless as Pandas dataframes (21) to connect important aspects of data acquisition through easy-to-follow abstractions. Finally, we aimed to be minimalistic from the perspective of package dependencies and from the perspective of re-using already existing tools in Python - for example, the said dataframes.

The first key OOP abstractions we have singled out are the *Experiment*, the *Specimen* and the *Phenotype*(fig. 1**A**). In this sense, the *Experiment* can be viewed as a logical container for several Petri dishes or multi-titre plates united by a similar experimental purpose. For example, an *Experiment* could be a virus titre determination performed in three biological replicas, which would combine three dishes into one OOP object. A *Specimen* in this example would constitute an individual dish or biological replicate. Within this dish, one may expect to find several plaques - i.e. *Phenotypes* in our abstraction. Combined, such a hierarchical structure is highly versatile and can cover a great variety of biological data. For example, irrespective of whether the experiment aims to perform the plaque assay by the means of an end-point Crytal Violet staining or use fluorescence microscopy, data can make use of the structure described above (fig. 1**B**).

**Fig. 1.**
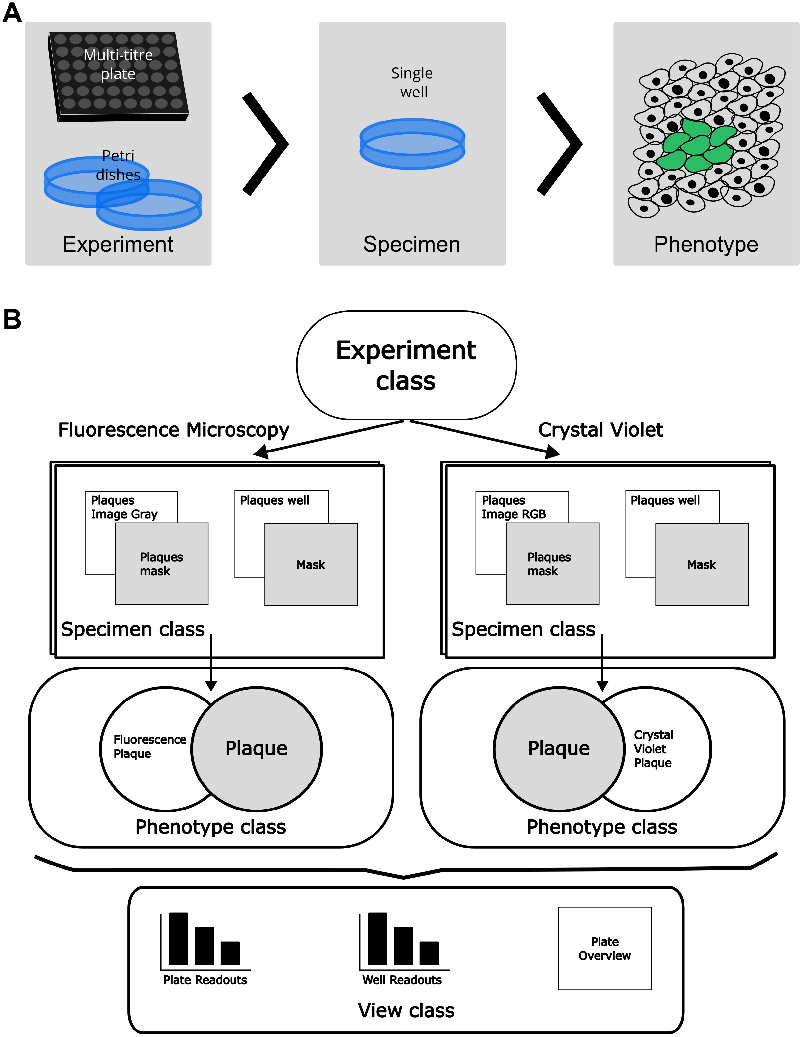
Object-oriented programming design of the PyPlaque package. *(A)* Schematic depiction of nested classes (grey rectangles with titles) representing the structure of PyPlaque abstractions accompanied by pictorial examples. *(B)* Flow diagram of the classes available to represent the experimental data. An example demonstrates how the Fluorescence Microscopy experiment or Crystal Violet experiment could make use of the same structures: *Experiment* > *Specimen* > *Phenotype*. A *View* module gives access to classes and methods for analysing different cross-sections of the experimental data for e.g., the whole plate, the well or the plaque.

The final key OOP abstraction we propose is the *View* classes. In our design, the *View* modules aim to contain any kind of OOP class aimed for analysis or visualisation obtained from the three classes mentioned above (see fig. 1**B**). The choice of the class to be used in order to construct a specific *View* defines the granularity of the results. For example, if the view is constructed for the granularity of the *Phenotype*, it would contain measurements of an individual plaque. Whereas if a *View* is constructed for the entire *Experiment* it would contain average measurements of plaques in this experiment. Once the classes are instantiated, this information is stored as nested dictionaries.

### Phenotypic Measurements Validation using Synthetic Data

Once plaque assay images are loaded into the respective data structures measurements can be performed using a respective *View*. Measurements can include count or morphological properties of the plaque phenotypes such as area or perimeter. However, different software platforms may perform such measurements differently, leading to a lack of comparability. To date the most popular platforms for biomedical image analysis include ImageJ/Fiji (19), Matlab (24) and Python (using packages like scikit-image (25)). Thus far, existing specialised packages for plaque quantification have been based on these platforms.

To understand how such measurements could be compared between these major platforms we have constructed a set of synthetic binary images depicting simple geometric shapes: an ellipse, a circle, two overlapping circles and a square (fig. 2), assigned to the numbers 1, 2, 3 and 4 respectively. These shapes were chosen to represent both regular and irregular shapes in the simplest way possible for ease of interpretation. To ensure comparability amongst the measurement systems, the shape parameters were identical while their location was chosen at random. fig. 2 shows 3 of 5 images used in the comparison.

**Fig. 2.**
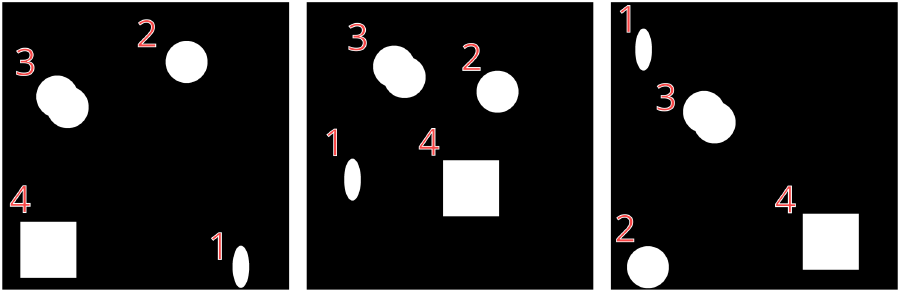
Analysis of Synthetic Images. Synthetic images generated with randomly located shapes of an ellipse, a circle, overlapping circles and a square with IDs, 1, 2, 3 and 4 respectively. 5 such images were generated and used for analysis and comparison between computation software (table 1, table 2) of which 3 are shown above.

Next, we performed two basic measurements of each object in the synthetic images using ImageJ/Fiji (19), Matlab (24) and our Python-based PyPlaque. Specifically, we have quantified the area and perimeter of these objects, as both of these measurements can be used to describe the extent of the virological plaques. In the case of PyPlaque, we used two different methods to estimate the area and perimeter. The first employed a built-in measurement using the scikit-image package (25). Second, was the method based on Pick’s theorem (see Page 9, **Pick’s Area and Perimeter**), which we developed specifically for PyPlaque.

**Table 1.** Object Area in Synthetic Images. Estimated values are averaged over 5 images where the objects were seeded in random locations. GT stands for ground truth. The performance closest to GT is marked in bold.

| Software | Area |  | ID |  |
| --- | --- | --- | --- | --- |
|  | ellipse | circle | overlapping circles | square |
| GT | 28274.334 | 70685.835 | 113733.351 | 160000.000 |
| PyPlaque (Pick’s) | 27588.000 | 69812.000 | 100779.000 | 158403.000 |
| Matlab (24) | **28233.000** | **70661.000** | **101806.000** | **160000.000** |
| ImageJ/Fiji (19) | **28233.000** | **70661.000** | **101806.000** | **160000.000** |
| PyPlaque (scikit-image) (25) | **28233.000** | **70661.000** | **101806.000** | **160000.000** |

**Table 2.** Object Perimeter in Synthetic Images. Estimated values are averaged over 5 images where the objects were seeded in random locations. GT stands for ground truth. The performance closest to GT is marked in bold. Second-best is underlined.

| Software | Perimeter |  | ID |  |
| --- | --- | --- | --- | --- |
|  | ellipse | circle | overlapping circles | square |
| GT | 690.384 | 942.478 | 1256.637 | 1600.000 |
| PyPlaque (Pick’s) | 704.000 | 988.000 | **1238.828** | <u>1604.000</u> |
| Matlab (24) | **686.456** | **939.000** | 1155.220 | 1563.716 |
| ImageJ/Fiji (19) | 722.943 | 991.561 | 1218.923 | **1597.657** |
| PyPlaque (scikit-image) (25) | 721.872 | 990.489 | <u>1219.024</u> | <u>1596.000</u> |

Area measurement results presented in table 1 suggest that all three platforms performed equally slightly underestimating the area compared to GT. Notably, while the PyPlaque area measurement using scikit-image (25) functionality performed equally to Matlab (24) and ImageJ/Fiji (19), the area measurement using Pick’s theorem led to even further underestimation.

Surprisingly, the perimeter measurement results presented in table 2 demonstrated an entirely different picture. In the case of *ellipse* and *circle* shapes, Matlab-based measurements were the closest to the GT with PyPlaque measurement based on Pick’s theorem showing the second-best results. In the case of the *square* shape ImageJ/Fiji demonstrated the best performance, with PyPlaque measurement based on scikit-image showing the second-best results. Finally, in the case of *overlapping circles*, PyPlaque measurement based on Pick’s theorem provided the measurement closest to the GT, with scikit-image performing second best. Crucially, Matlab, Image/Fiji and scikit-image measurements underestimated the perimeter severely on the overlapping circles.

Together our results suggest a surprising incomparability of simple shape measurements across modern bioimage analysis platforms. However, due to the flexibility coming with using a Python package, the user is free to choose the measurement approach depending on the needs of the project. It is worth noting that we have additionally performed the area and perimeter measurements with another popular Python library OpenCV (26). The results of these measurements were largely identical to scikit-image and therefore omitted here.

### Crystal Violet Plaque Assay Measurements at varying Granularity using PyPlaque

To demonstrate how the structure depicted in fig. 1 can be used to store and analyse data from a conventional plaque assay we performed an analysis of a conventional assay plaque of Vaccinia Virus (VACV) stained with Crystal Violet. The images of the assay plates were taken from an open dataset available online (27). This dataset consists of mobile photographs of the plaque assay plates accompanied by binary masks for wells and individual plaques. Each image of the dataset contains objects of increasing granularity containing quantitative information about the assay.

Specifically, fig. 3**A**, shows a 6-well plate (left) and a masked version of the full 6-well plate (right). Such data can be handled by the ***PlateImage*** class of PyPlaque. fig. 3**B** goes further in granularity showing a single well from the plate (left) and the corresponding masked version of the well (centre). Such data is handled by our classes ***PlaquesWell***. The rightmost column of fig. 3**B** inset is a single viral plaque from the well. We have the functionality using our class ***PlaquesMask*** to extract readouts such as the average size of all the plaques in the well, the total count of plaques and the area of an individual plaque as also shown in the figure.

**Fig. 3.**
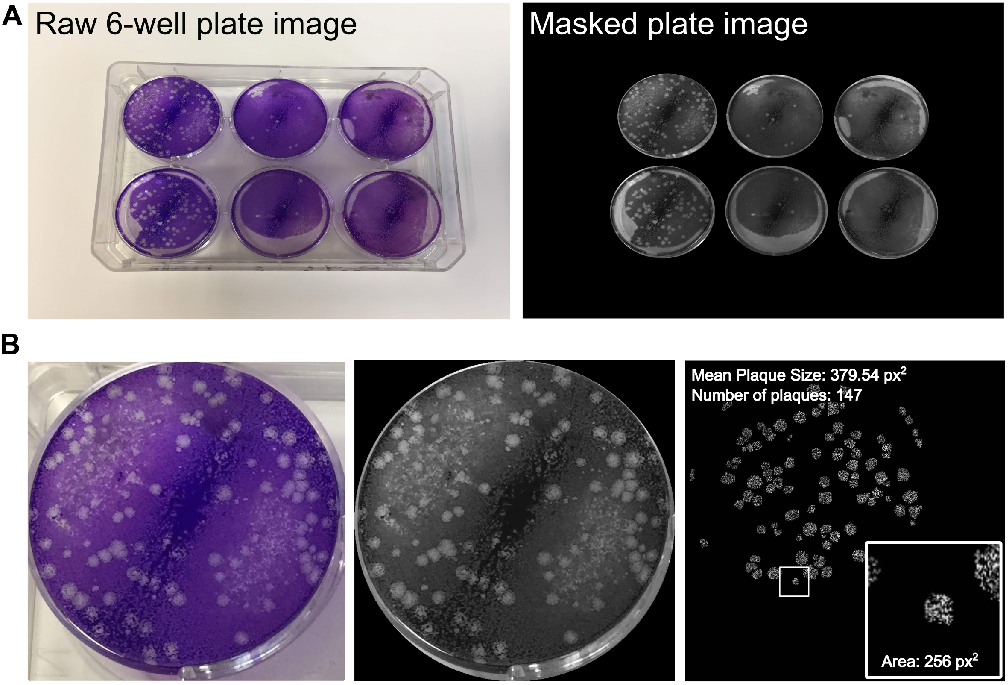
Granularity in the Detection of Objects in Crystal Violet Plaque Assay Images of VACV. Objects with increasing granularity are stored and analysed by PyPlaque. *(A)* Shows a 6-well plate *(left)* and a masked version of the full 6-well plate *(right). (B)* Shows a single well from the plate *(left)*, corresponding masked version of the well *(centre)* and a single viral plaque from the well *(right-inset)* with average size of the plaques in the well, the number of plaques *(top-left)* and area of an individual plaque *(inset bottom-right)*.

The usage of the classes mentioned above are further explained in the [crystal violet plaque](https://github.com/plaque2/PyPlaque/blob/master/notebooks/cvp_detailed.ipynb) notebook found in the [GitHub](https://github.com/plaque2/PyPlaque) page of the project.

The granularity provided by PyPlaque readouts could be beneficial in comparing plaques across wells treated with virus inoculum at different dilutions. Furthermore, it may allow comparing measurements across plates and different experiments.

### Scalable Plaque Analysis using PyPlaque in High– content Fluorescence Microscopy

Beyond measuring the titre in the virus inoculum, plaque assay can be employed to measure virus spread under conditions of perturbation. Techniques like high-content fluorescence microscopy allow scaling up such efforts (11, 28). To illustrate how PyPlaque can be employed in quantifying such experiments we have performed an analysis of a control 384-well plate half of which was infected with GFP-transgenic human coronavirus OC43 (CoV-GFP). This plate was obtained from a published dataset (29). The control well plate we used is imaged for two fluorescence channels: CoV-GFP and Nuclei (Hoechst 33342 dye).

Comparably to the PyPlaque workflow constructed for the Crystal Violet plaques, information in the high-content fluorescence microscopy exists at several levels of granularity fig. 4**A**. In contrast to the Crystal Violet plates, however, individual images in fluorescent microscopy are obtained at the level of wells. That means that the lower granularity plate overview is not available and must be constructed dynamically. This can be achieved using the ***View*** class. Results in the fig. 4**A** show that, as expected, the left-hand half of the control plate has visible GFP-positive cells, some of which are clustered into fluorescent CoV plaques. Notably, some wells on the right-hand side contain sample preparation artefacts (e.g. dust and fibres) (30), which may contribute to the background detection due to autofluorescence (fig. 4**A**, zoomed insets).

**Fig. 4.**
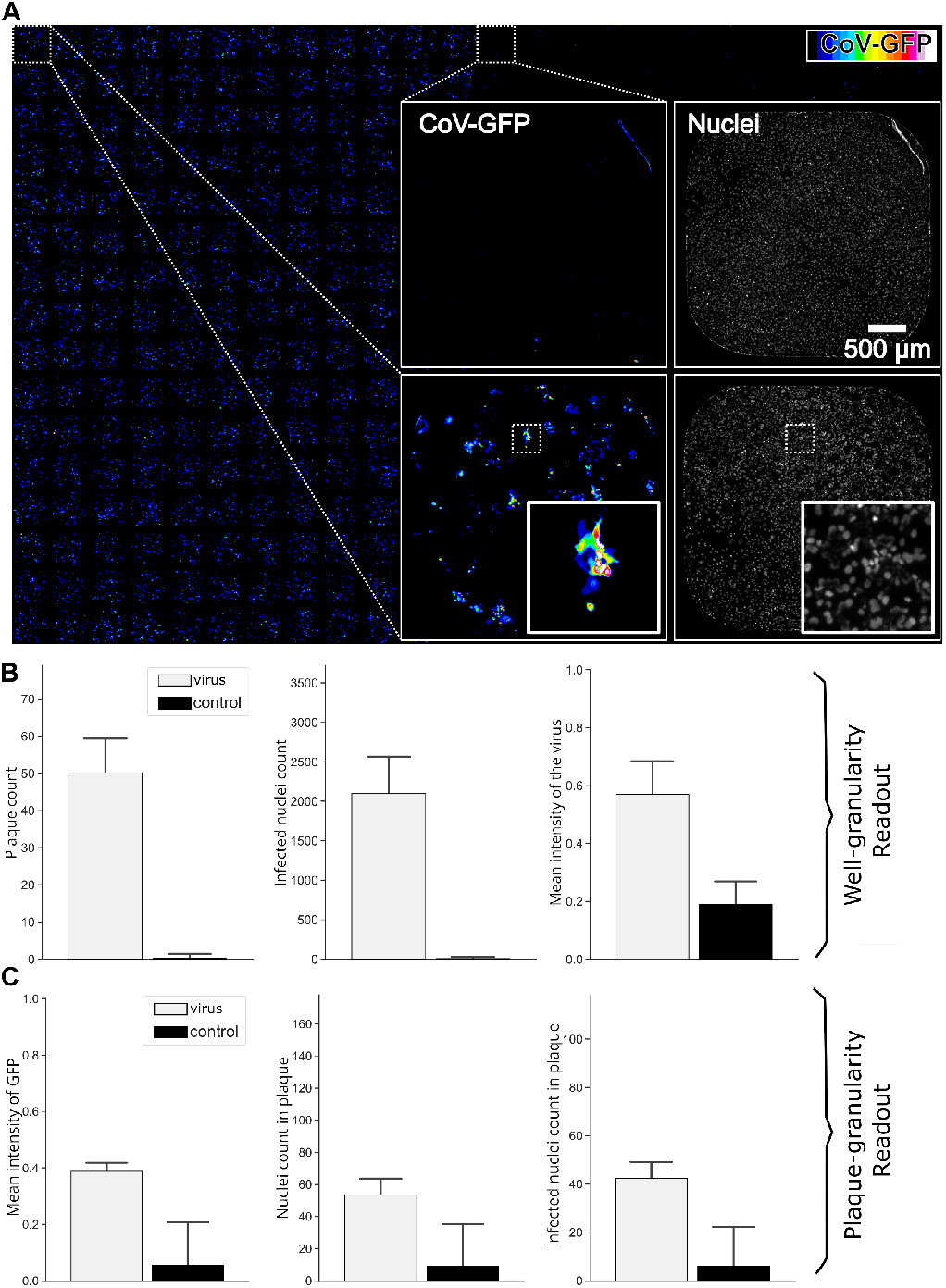
Pyplaque measurements at image and object level. PyPlaque (20) includes methods to provide readouts at an image and object level. *(A)* Example 384-well Fluorescence microscopy imaged plate. Insets show examples of two wells from two vertical halves of the entire 384-well plate. Left column of the insets show the CoV-GFP channel and right column shows the Nuclei (Hoechst dye) channel. Insets within the insets show a section of a single well for the CoV-GFP and Nuclei channel on the left and right respectively. *(B)* Plot of readout values at the well level from the software for the plate shown in panel (A). *(C)* Plot of readout values at the object level for the plate, averaged first over wells. For each plot, we have the infected wells *(left)* and control wells *(right)* indicated as virus and control in the legend. Such readouts could be generated by the classes and methods from *View* (see fig. 1).

Noteworthy, fluorescence microscopy does not usually come with segmentation masks. While segmentation masks can be obtained through advanced deep learning models, in some cases rule-based algorithms like Otsu (31) are sufficient. Both cases are supported by PyPlaque. However, while rule-based thresholding and connectivity handling is built-in via ***get_all_plaque_regions*** method, the former requires an external model.

Results of quantifications using built-in methods can be seen in fig. 4**B** and **C**. Specifically, fig. 4**B** demonstrates differences between the infected and uninfected halves of the plate (denoted as virus and control respectively) at the granularity of a well. Measurements like plaque count, infected nuclei count and the mean intensity of the virus signal can be obtained using the built-in methods of the PyPlaque package. fig. 4**C** demonstrate that such analysis is also possible at the level of individual plaque. Measurements such as nuclei count in the plaque, and infected nuclei count in the plaque, as well as mean GFP intensity can be obtained. The latter, in turn, could facilitate the separation of real plaque from a sample preparation artefact by simply filtering out the objects with very low fluorescence during the post-processing of the analysis results. For full list of measurements, please see Page 8, **Classes for well and plaque level readouts**.

Importantly, workflows for the data source as different as mobile photography and high-content fluorescence microscopy can be flexibly constructed using the OOP structure we propose for PyPlaque (20). In the spirit of Python Data Science, these workflows can be implemented as Jupyter (23) notebooks. An example of such an [analysis notebook](https://github.com/plaque2/PyPlaque/blob/master/notebooks/visualization/example_analysis.ipynb) can be found in the [GitHub](https://github.com/plaque2/PyPlaque) page of the project.

## Discussion

Quantification of phenotypes in the virological assays plays an important role in Virology research and our search for novel antivirals (32, 33). In the case of the virological plaque assay, several quantification tools have been proposed in the past (9, 11, 13–18). However, the overall focus on the graphical user interface (GUI) makes these tools unflexible and hard to maintain. In many cases, the GUI part of the software is as complex as all the other functionalities (11). Furthermore, GUI shifts the focus of the package, forcing a particular workflow design.

The PyPlaque (20) package developed in this work takes a different approach to the quantification of virological assays. Rather than providing a GUI-based solution, we propose a Python package rooted in the Data Science technological stack. Instead of forcing the user to follow a particular workflow, we provide an abstract OOP container for experimental data that can be integrated into Pythonic workflow constructors like Jupyter (22, 23). We argue that by making the data constructs available in the Jupyter environment, our approach is more scalable and flexible due its to compatibility with modern Data Science tools like Pandas (21). In this sense, once the data is ingested, running the analysis is as simple as putting an action into a for-loop - simplicity accessible to users from various backgrounds.

Incidentally, this approach allowed us to focus on the actual measurements providing users with the choice of methods. In a series of comparisons we demonstrated the lack of agreement in a seemingly trivial task of perimeter measurement in the leading BioImage analysis platforms including Matlab (24), ImageJ/Fiji (19) and Python scikit-image (25). Notably, the go-to Python approach showed the greatest disagreement. To remedy this, we propose an alternative perimeter measurement approach alleviating the disagreement in certain cases.

Furthermore, we argue that the approach proposed here can be generalised to other biological assays yielding a new paradigm. Similar structures of classes at varying granularities from ***Experiment*** to ***Phenotype*** can be constructed for many kinds of experiments. In the particular case of experiments conducted with sample carriers shown here, PyPlaque can be used for assays other than plaque assay already. We hope to receive the open-source community contributions to further expand this scope.

## ACKNOWLEDGEMENTS

This work was partially funded by the Center for Advanced Systems Understanding (CASUS) which is financed by Germany’s Federal Ministry of Education and Research (BMBF) and by the Saxon Ministry for Science, Culture, and Tourism (SMWK) with tax funds on the basis of the budget approved by the Saxon State Parliament. We thank Urs Greber (University of Zurich, Switzerland) for access to raw data of (29). The authors acknowledge the financial support by the Federal Ministry of Education and Research of Germany and by Sächsische Staatsministerium für Wissenschaft, Kultur und Tourismus in the programme Center of Excellence for AI-research “Center for Scalable Data Analytics and Artificial Intelligence Dresden/Leipzig”, project identification number: ScaDS.AI.

## Methods

### PyPlaque Installation

The release version of the PyPlaque package can be installed using PyPi package manager by invoking the following command in the command line interface:

- ***pip install pyplaque*** The developer version can be installed directly from the *master* and *dev* branches of the [GitHub](https://github.com/plaque2/PyPlaque) repository.

### Multi-granularity Level Design

As mentioned above the design of the PyPlaque classes follows a multi-level granularity approach. This means that the structures of the package allow to group data and metadata at different levels, preserving semantic relationships. These structures are detailed in the subsequent sections.

### Experiment Granularity Level Classes

At the highest level of granularity, we defined classes ***CrystalViolet*** and ***FluorescenceMicroscopy*** which are designed to contain metadata of multiple instances of a multi-titre plate of Crystal Violet Plaque and Fluorescence Plaque respectively. To aid in having a better overview of all of the multi-titre plates in the experiment, both of these classes contain methods ***get_individual_plates, get_number_of_plates*** and ***read_from_path*** that identify the individual plates, their total count and read each individual well image respectively.

### Single Plate Granularity Level Classes

At the next level of granularity are the classes aimed at containing data and metadata for one experimental container, for example corresponding to a single multi-titre plate. The ***PlateImage*** class accepts an image and corresponding binary masks of individual wells of a single full multi-titre plate. Using class properties such as the number of rows, number of columns and a flag indicating whether or not the plate was imaged inverted or not, given by the user, methods of the class return the following:

- ***get_wells*** - returns a list of the images of individual wells of the plate stored as binary Numpy arrays.
- ***get_well_positions*** - returns a dictionary where individual wells of the plate are the units. For each individual well, the image, mask, masked image, bounding box x-y values, row and column number estimated from the well positions are stored with the rows and columns numbered starting from 0.
- ***plot_well_positions*** - returns a plot with bounding boxes drawn around individual wells of the plate (inferred from the plate mask), with rows and columns numbered starting from 0.

### Single Well Granularity Level Classes

The class ***PlaquesWell*** is aimed at containing an individual well of a multi-titre plate. It expects information about the row and column numbers, the well image and mask. It has the method ***get_image*** which returns a masked image of the individual well.

The ***PlaquesMask*** is a feature-enriched class designed with the aim of holding a single binary mask containing multiple plaque instances from an individual well. Using the binary mask of plaques and a name to identify this plaque mask (usually derived from the plate name and the well position that the mask corresponds to), methods of the class return the following:

- ***get_plaques*** - returns a list of individual plaques stored as binary Numpy arrays.
- ***get_measure*** - returns measures based on the individual plaques in a well as a dictionary. This is more appropriate for cumulative measurements as compared to the ***measure*** method under class ***Plaque*** (see Page 6) that gives granular measurements based on each plaque.
- ***plot_centroid*** - plots a dotted ring around all the plaques that are found in a well. This ring is centred at the centroid (see Page 8, ***get_centroid***) of all the centres of individual plaques found.

The classes ***PlaquesImageRGB*** and ***PlaquesImageGray*** are child classes of ***PlaquesMask***. These two classes are designed to hold grayscale image data and RGB image data containing multiple plaque instances respectively with a respective binary mask. At first glance, these classes could be thought of as the same with a flag differentiating the channel in the image they can hold. However they are primarily different because of the methods that they give the user access to while opting for one or the other. Similar to the class ***PlaquesMask***, these two classes too use a name (usually derived from the plate name and the well position that the mask corresponds to) to identify the plaque image and mask object it contains. ***PlaquesImageGray*** additionally contains accepts two parameters *threshold* and *sigma* which are further used to apply Gaussian blur and binarise the grayscale image.

### Single Phenotype Granularity Level Classes

The class ***Plaque*** is designed to contain a single virological plaque phenotype as an object. It encapsulates the properties related to a specific plaque, including its mask, centroid coordinates and bounding box, and offers methods for the calculation of the area, eccentricity and roundness of this plaque object. Methods of the class return the following:

- ***measure*** - returns the bounding box area (in pixels) of a plaque object, as well as an approximation of its actual area based on the proportion of white pixels in the mask.
- ***eccentricity*** - calculates and returns the eccentricity (34) of an individual plaque object. The eccentricity is determined by fitting an ellipse to the plaque’s boundary contour and using the formula,

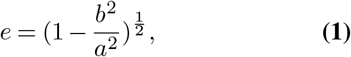

where *b* represents the length of the semi-minor axis, and *a* represents the length of the semi-major axis.
- ***roundness*** - returns the roundness (35) of an individual plaque object. The roundness is calculated using the formula,

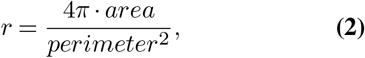

where *area* is estimated as *πR*^2^, and *perimeter* is calculated as 2*πR, R* being the radius estimated from the centre to the top-right corner of the bounding box. This method can alternatively use Pick’s (see Page 9, **Pick’s Area and Perimeter**) area and perimeter estimates. This value is 1 in the case of a perfect circle.

The classes ***CrystalVioletPlaque*** and ***FluorescencePlaque*** are child classes of the class ***Plaque***. They allow for the storage of data and metadata of crystal violet or fluorescent dye stained plaques imaged using mobile photography or bright-field microscopy and fluorescence microscopy respectively. Since the masks are binary in both cases, both of these classes are able to use the methods such as ***measure, eccentricity***, and ***roundness*** from the parent class.

### Analysis of Nuclei

The analysis of nuclei begins with the acceptance of a well image corresponding to the nuclei channel stained with fluorescent dye and imaged via fluorescence microscopy. In case a pre-generated mask is available it is also read simultaneously with the images into the plate dictionary structure provided by our method ***load_wells_for_plate_nuclei()*** available under the class ***FluorescenceMicroscopy***. In case a mask isn’t present, it is generated at runtime, using the following steps, some of which are implemented in Python in the package OpenCV (26).

1. **Artefact Removal (based on Intensity)** - *X*_*i,j*_ = 0 if *X*_*i,j*_ *> α* else *X*_*i,j*_, where *α* is the ‘artefact_threshold’ for the nuclei channel.
2. **Background Noise Removal (based on Structure)** -
  a. Dilation *X*′ = *X* ⊕ *B* =⋃_*b*ϵ*B*_ *X*_*b*_, where *X*_*b*_ is the translation of *X* by *b*. To be more specific,

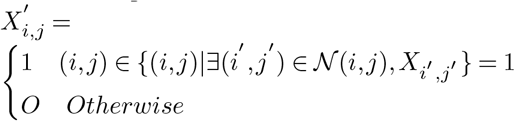
  b. Erosion 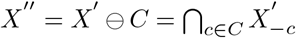, where 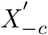 is the translation of *X*′ by −*c*. To be more specific,

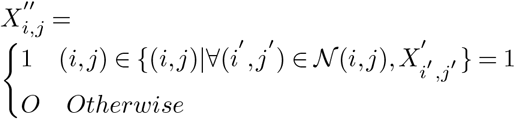

when we choose *B* as a sub-matrix of *C* it leads to the removal of the background noise *B* and *C* are chosen for the matrix representation of a circle of diameter, *d* = 2× ‘correction_ball_radius’ for the nuclei channel.
3. **Binarise** – 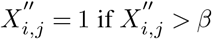 else 0, where *β* is the ‘manual_threshold’ for the nuclei channel.

For readouts relating to the nuclei channel please look at the Page 8, **Classes for well and plaque level readouts**.

### Analysis of Virus Signal

The analysis of the virus signal begins similarly to nuclei with the acceptance of a well image corresponding to the virus channel stained with crystal violet or fluorescent dye and imaged via brightfield or fluorescence microscopy. In case that a pre-generated mask is available it is also read simultaneously with the images into the plate dictionary structure provided by our method ***load_wells_for_plate_nuclei()*** available under the class ***FluorescenceMicroscopy*** or ***load_well_images_and_masks_for_plate()*** available under the class ***CrystalViolet***. The relevant notebooks in the GitHub repository contain practical examples.

In case a mask isn’t present, it is generated at runtime just like nuclei. But since there is a difference in how virus channel appears in crystal violet or fluorescent dye stained samples and also under mobile photography, bright-field or fluorescence microscopy, we have two different workflows for mask generation for plaques. Once again, part of these workflows are implemented in Python in the packages OpenCV (26), Scikit-Image (25) and SciPy (36). The workflows are as follows:

### Virus Mask Generation for Fluorescence Microscopy

1. **Binarise** - *X*_*i,j*_ = 1 if *X*_*i,j*_ *> β* else 0, where *β* is the ‘virus_threshold’ for the virus channel.
2. **Connected Components** – 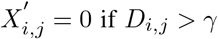 else 1, where *D*_*i,j*_ is the distance to the nearest foreground object and *γ* is the ‘plaque_connectivity’ for the virus channel.

### Virus Mask Generation for Mobile Photography or Brightfield Microscopy

1. **Gaussian Blurring** - 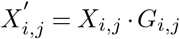, where 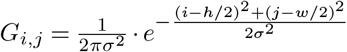 where (*w, h*) is the shape of *X* and *σ* is the ‘sigma’ passed to ***PlaquesImageGray*** Specimen class.
2. **Binarise** - 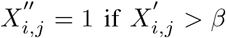 else 0, where *β* is the ‘threshold’ passed to ***PlaquesImageGray*** Specimen class.

For readouts relating to the virus channel please look at Page 8, **Classes for Well and Plaque Level Readouts**.

### Classes for Well and Plaque Level Readouts

PyPlaque returns readouts at the well and plaque level. It is worth mentioning that the modules described in this subsection are intended for and designed to work for fluorescence microscopy images. In this section, we discuss the classes that are used for such analysis and proceed to describe how we compute each of the readouts including those that are showcased in Fig. fig. 4**B** **and C**.

The classes ***PlateReadout, WellImageReadout*** and ***PlaqueObjectReadout*** can be used to deal with the analysis at different granularity levels. They are used to generate readouts pertaining to multiple wells of a single plate of fluorescence plaques, multiple instances of plaques within a single well and finally a single instance of a plaque from a fluorescence plaque well respectively. It is possible to obtain readouts at the well and/or plaque level based on flags passed to ***PlateReadout***.

Examples of readouts at the well level obtained by methods under ***WellImageReadout*** are:

- ***get_max_plaque_intensity*** - returns *max*(*X*_*v*_), where *X*_*v*_ is the image from the virus channel for the well.
- ***get_mean_plaque_intensity*** – returns 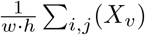, where *X*_*v*_ is as above with shape (*w, h*).
- ***get_total_plaque_intensity*** - returnsΣ _*i,j*_ (*X*_*v*_), where *X*_*v*_ is as above.
- ***get_median_plaque_intensity*** - returns *q* s.t. 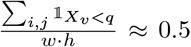, where *X*_*v*_ is as above and 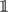 is the indicator function taking value 1 when the condition is satisfied and zero otherwise.
- ***get_nuclei_count*** - returns 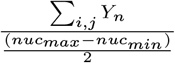 where *Y*_*n*_ is the mask from the nuclei channel for the well and (*nuc*_*min*_, *nuc*_*max*_) are the upper and lower bounds for the area of a nucleus.
- ***get_plaque_count*** - returns the length *l* of the list of global maxima coordinates identified from *X*_*v*_ which is as described above. It is an extension of the steps mentioned in Page 7, ***Virus Mask Generation for Fluorescence Microscopy*** the further steps being,

3. **Individual Plaques** - getting **regionprops** (from scikit-image (25)) objects based on Connected Components (37) that are greater than *vir*_*min*_, the lower bound for the area of a plaque from an image like 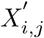 mentioned in Page 7, textit**Virus Mask Generation for Fluorescence Microscopy**.
4. **Counting global maxima in blurred masked plaque image** - For each plaque object, 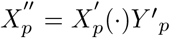 where 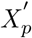 and 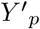 are the image and mask of a single plaque *p* respectively. Then we use Gaussian blurring with a specified *σ* as described in Page 7, ***Virus Mask Generation for Mobile Photography or Brightfield Microscopy*** to get 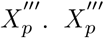 is then used to identify global maxima (from scikit-image (25)) within it that are identified based on the hyper-parameter *min*_*d*_ so that the distance between two global maxima found is at least (2 *· min*_*d*_ + 1) indicating that the spread of the area for which a global maxima is found is at least of radius *min*_*d*_. This operation dilates (see Page 7, **Analysis of Nuclei**) and merges neighbouring maxima that become part of the same connected component after dilation. Coordinates (*x, y*) of the locations where 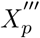 is equal to the dilated image are returned as global maxima. The length of this list of coordinates can therefore be measured.
- ***get_infected_nuclei_count*** – returns 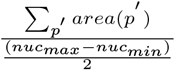 where *p*′ ∈{plaques identified from *Y*_*v*_ using step 3 under ***get_plaque_count*** mentioned previously}, *Y*_*v*_ being the mask of the virus channel for the well, (*nuc*_*min*_, *nuc*_*max*_) are as described previously and *area*(*p* ′.) is simply the area of the plaque *p* ′.
- ***get_lesion_area*** – returns Σ _*i,j*_ (*Y*_*v*_), where *Y*_*v*_ is as mentioned above.
- ***get_plaque_objects*** - returns the **regionprops** (from scikit-image (25)) objects identified from *Y*_*v*_, where *Y*_*v*_ is as mentioned above, using step 3 under ***get_plaque_count*** mentioned above.

Other than the methods mentioned above, the class provides the following methods such as ***get_nuclei_image_name, get_plaque_image_name, get_row, get_column, get_max_nuclei_intensity, get_mean_nuclei_intensity, get_total_nuclei_intensity, get_median_nuclei_intensity*** as well. They have not been described in detail since they are either similar to the ones described and can be thought of analogously or they are self-explanatory from their names.

The class calls via the ***call_plaque_object_readout*** method the readouts at the plaque level. An example of readouts at the plaque level obtained by methods under ***PlaqueObjectReadout*** are:

- ***get_area*** - returns **regionprops** (from scikit-image (25)) area of the plaque, Σ_*i,j*_ *Y*_*p*_, where *Y*_*p*_ is the mask of the plaque or if indicated by a flag the Pick’s area (see Page 9, **Pick’s Area and Perimeter**) based on *Y*_*p*_ is returned.
- ***get_perimeter*** - returns **regionprops** perimeter of the plaque, or if indicated by a flag the Pick’s perimeter (see Page 9, **Pick’s Area and Perimeter**) based on *Y*_*p*_ is returned.
- ***get_centroid*** - returns **regionprops** centroid of the plaque, 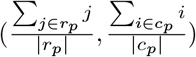 where |*r*_*p*_ | and |*c*_*p*_| are the number of pixel rows and columns of the plaque mask *Y*_*p*_ or the plaque image *X*_*p*_.
- ***get_bbox*** - returns a tuple (*x, y, w*_*p*_, *h*_*p*_) where (*x, y*) are the bounding box coordinates from the top left corner of the plaque and (*w*_*p*_, *h*_*p*_) is the shape of bounding box of the plaque.
- ***get_major_minor_axis_length*** - returns a tuple (4*a*, 4*b*) where *a* and *b* are as described in Eq. (1).
- ***get_eccentricity*** - returns the value of eccentricity as described in Eq. (1).
- ***get_roundness*** - returns the value of roundness (35) as described in Eq. (2).
- ***get_number_of_peaks*** - returns the length of a list of coordinates (*x, y*) of the global maxima found in the masked image of the plaque (similar to 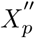) using the method mentioned in step 4, **Counting global maxima in blurred masked plaque image** of ***get_plaque_count*** under the well level readouts.
- ***get_nuclei_in_plaque*** – returns 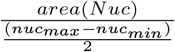 where (*nuc*_*min*_, *nuc*_*max*_) are as described previously and *area*(*N uc*) is either the sum of non-zero pixels or if indicated by a flag, the Pick’s area (see Page 9, **Pick’s Area and Perimeter**) of a masked image *N uc* defined as 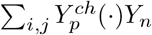 where 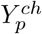 is the **regionprops** convex-hull (38) of the plaque mask *Y*_*p*_ and *Y*_*n*_ is the corresponding nuclei mask from the nuclei channel.
- ***get_infected_nuclei_in_plaque*** – 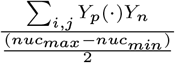 where (*nuc*_*min*_, *nuc*_*max*_), *Y*_*p*_ and *Y*_*n*_ are all as described previously.
- ***get_convex_area*** - returns **regionprops** convex area of the plaque which is the sum of non-zero pixels in a binary image of the convex hull (38) of the plaque.
- ***max_GFP_intensity*** - *max*(*X*_*p*_(*i, j*(*·*)*Y*_*p*_(*i, j*)), where *X*_*p*_ and *Y*_*p*_ are the image and mask of the plaque *p*.
- ***total_GFP_intensity*** -Σ _*i,j*_ *X*_*p*_(*i, j*)(*·*)*Y*_*p*_(*i, j*), where *X*_*p*_ and *Y*_*p*_ is as above.
- ***mean_GFP_intensity*** – 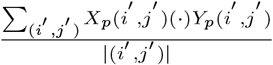 where *X*_*p*_ and *Y*_*p*_ is as above and (*i*′, *j*′)represents a non-zero pixel in *X*_*p*_(*·*)*Y*_*p*_.

Other than the methods mentioned above, the class provides the following methods such as ***get_row, get_column*** as well. For getting the means, maximums and totals and storing readouts in dataframes, implementations available from Numpy (39) and Pandas (21) are used. The readouts for at the plaque level are averaged for all plaques within a well before being stored in a dataframe but individual plaque measurements can be accessed as shown in our [fluorescence plaque detailed](https://github.com/plaque2/PyPlaque/blob/master/notebooks/fp_detailed.ipynb) notebook meant for advanced users of the software.

Further details of the readouts at both levels can be found in the [view](https://github.com/plaque2/PyPlaque/tree/master/PyPlaque/view) module in the [GitHub](https://github.com/plaque2/PyPlaque) page of the project.

### Pick’s Area and Perimeter

The Pick’s Area and Perimeter used here is the main result of Pick’s theorem (40), first showcased by Georg Alexander Pick. It is a geometric result for calculating the area and perimeter of irregular polygons on integer lattices (41). The theorem states that given a simple lattice polygon ***P***, if ***i*** is the number of integer lattice points inside it and ***p*** is the number of integer lattice points on its perimeter, then the area of the polygon ***area(P)***, is given by,

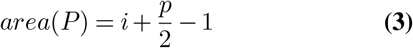

where a point (*x, y*) ∈ℝ^2^ is an integer lattice point, or simply lattice point, ***iff*** both *x* and *y* are integers. We define a lattice polygon to be a polygon all of whose vertices are at integer lattice points.

It is important to note that the same formula does not apply for polygons with holes where it changes to,

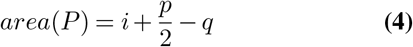

where *q* depends on the number of holes. For a proof of the theorem please look at the work here (42).

As for the perimeter and area calculations in our code, we calculate the perimeter in both cases and use it along with the interior measurements to get the total area and simply return the perimeter estimate for the perimeter. We can do so using the following steps:

1. Creating a border image *X*′ using Dilation or Erosion (see Page 7, **Analysis of Nuclei**) and subtracting or subtracting it from the original respectively. The eroded image forms the interior of the polygon. The neighbourhood definition *N* (*i, j*) defines the result of these operations. For our purposes,

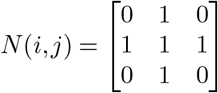
2. We embolden the resulting border image by convolution (43), *X* ″=*X* ′* *K* where,

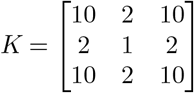
3. Taking a weighted average of the histogram of the above-mentioned convolved image *X* ″, giving appropriate weightage to the intensity values in the bins of the histogram. For computational simplicity, we keep the number of bins at 50.

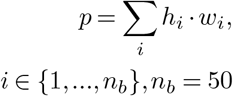

